# Desensitization of opsin responses during all-optical interrogation depends on imaging parameters

**DOI:** 10.1101/2025.10.02.680058

**Authors:** Blake Russell, Robert M. Lees, Armin Lak, Adam M. Packer

## Abstract

**Significance:** The combination of two-photon calcium imaging and two-photon optogenetic stimulation, termed all-optical interrogation, provides spatial and temporal precision when recording and manipulating neural circuit activity in vivo. All-optical experiments often use red-shifted opsins in combination with green fluorescent reporters of neuronal activity. However, their excitation spectra still partially overlap, meaning that the imaging laser can excite the opsin. Though some care has been taken in the past to understand the effects of this spectral overlap, further work is required to understand its impact on the findings of all-optical studies.

**Aim:** We aimed to investigate whether two-photon imaging of the green fluorescent calcium reporter GCaMP6s at 920 nm increases the rate of desensitization in neurons expressing the red-shifted opsin C1V1.

**Approach:** We systematically varied either the inter-stimulus interval or the duration of two-photon calcium imaging during two-photon optogenetic stimulation of mouse layer 2/3 barrel cortex or visual cortex neurons.

**Results:** We found that two-photon imaging at 920 nm increases the desensitization of photostimulation responses across trials in C1V1-expressing neurons - an effect that is exacerbated at shorter inter-stimulus intervals. Reduced photostimulation responses are not limited to targeted cells, and are found across the entire field of view. Such network effects are less pronounced at shorter imaging doses.

**Conclusions:** Our results provide methodological optimizations that enable opsin desensitization to be mitigated in all-optical experiments. This will reduce an external source of trial-by-trial variability in future all-optical experiments.

## 1. Introduction

All-optical interrogation combines imaging of fluorescent reporters of neuronal activity with optogenetic stimulation to simultaneously measure and manipulate neuronal activity in vivo^1,2^. The most widely used optogenetic methodology utilizes channelrhodopsins, light-gated ion channels expressed in algae^3^, to optically control the activity of populations of neurons that can be defined by their genetics, using specific promoters to express the channelrhodopsin^4^, or through spatially-confined viral injection into a specific brain region^5^. Further, single-cell resolution of optogenetic stimulation can be achieved using two-photon excitation^6,7^, due to the probability of excitation being confined to a small volume^8^. Directing the stimulation laser through a spatial light modulator (SLM)^9^ allows holographic photostimulation of tens of neurons simultaneously^2^ and hundreds of neurons pseudo-simultaneously by rapidly switching the hologram^10^. Crucially, concurrent activation of tens of cortical neurons is enough to modulate existing behaviors^11–14^ and even drive new behaviors^10,14–16^. Using an SLM during all-optical interrogation enables the recreation of cortical population activity generated by externally-presented stimuli. This positions all-optical experiments with a unique advantage in exploring causal relationships between neuronal activity and behavior^17^. However, in order to artificially generate endogenous-like firing patterns, the power and scan pattern of the photostimulation laser has to be carefully calibrated using electrophysiology^2,18^. Therefore, any variable that can affect an opsin’s stimulus-evoked photocurrent can have a potentially large impact on the validity of all-optical studies. One such variable is opsin desensitization: the decrease in opsin activation due to repeated light exposure.

Like ‘bleaching’ in their G-protein-coupled counterparts^19–21^, channelrhodopsins desensitize under continuous stimulation and recover during darkness^22^. Even when not exposed to continuous stimulation, their rapidly peaking initial transient response to activation decreases exponentially across successive stimulations. The mechanism by which channelrhodopsins desensitize has been extensively characterized^22–29^. Opsin desensitization is distinct from the adaptation mechanisms that generate a decreasing response of cortical sensory neurons to repetitive naturalistic stimulation, which are primarily driven by changes in subcortical input instead of intrinsic properties and can be overcome with direct stimulation^30^. Additionally, the decrease in response caused by opsin desensitization is more severe than adaptation^31^, and as such, reduced responses to successive photostimulations of target cells are likely dominated by opsin desensitization.

Since 2010, our understanding of the mechanisms behind opsin desensitization have largely gone unchanged, settling on a two-photocycle model where ground and desensitized photocycles are concurrently populated with different efficiencies by a population of opsin molecules^29^. Because transitioning from the desensitized photocycle to the ground photocycle is ∼30x slower than the reverse scenario^23^, inter-stimulus intervals shorter than the time needed for all molecules to resensitize results in the accumulation of molecules in the desensitized photocycle. This can be circumvented by changing the identity of stimulated neurons between trials, an advantage more easily conferred by holographic-stimulation compared to one-photon bulk activation^2^. In an all-optical setup, changing the identity of stimulated neurons by swapping the SLM phase-mask would theoretically mitigate opsin desensitization whilst keeping the imaging field of view constant. However, the contribution of persistent imaging laser excitation to opsin desensitization remains critically understudied.

With simultaneous reading and writing of neural activity comes the desire for spectrally distinct opsins and fluorescent reporters. The issue of cross-talk, where the excitation wavelength used for the reporter also excites the opsin, has long been an issue with all-optical studies. In an attempt to alleviate this issue, many developments have been made to try and strike a balance between spectral distinction and optimal sensor/actuator dynamics. However, the most popular combination for all-optical interrogation remains the ‘original choice’: GCaMP as the sensor and C1V1 as the opsin. The GCaMP series of genetically encoded fluorescent proteins are optimal for calcium imaging in superficial layers of the cortex, having been iteratively improved upon and engineered to enhance signal-to-noise ratio and accelerate kinetics over the past two decades^32^. Because GCaMP is excited by blue light^32^, widely used blue-shifted opsins such as channelrhodopsins 1 and 2^3,22^ are not optimal for all-optical studies. Instead, the original choice of opsin for all-optical interrogation was C1V1^1,2,33,34^(but see^35^), a red-shifted variant of Channelrhodopsin-1^36^. C1V1 has previously been shown to desensitize over trials in an all-optical paradigm in vivo with a 10 second inter-stimulus interval^14^ and requires ∼30 seconds to fully resensitize under one-photon stimulation in vitro^37^. Because C1V1 is red-shifted, it has a high absorption cross-section at the near infrared wavelengths utilized for two-photon excitation^38^, with the greatest photocurrents occurring at 1040 nm^39^. Although 1040 nm is far from the peak of GCaMP’s two-photon excitation spectra (both calcium-bound and unbound), C1V1 still has a significant cross-section at 920 nm - GCaMP’s optimal two-photon excitation wavelength^32^. It has previously been shown that two-photon excitation at ∼900 nm causes an increase in the photocurrents of red-shifted opsins^40,41^, including C1V1^1^. Specifically, firing rates of C1V1-expressing neurons increase with increasing laser dwell time per cell^2^ and power on sample^34^. However, it remains unknown whether imaging parameters that are subthreshold for significantly increasing firing rates of C1V1-expressing neurons are able to transition opsin molecules into the desensitized photocycle and thus affect response dynamics.

In this study, we aimed to investigate whether the rate of desensitization in neurons expressing the opsin C1V1 increases when imaging GCaMP6s at 920 nm with common experimental two-photon imaging parameters that are subthreshold for eliciting firing alone. If so, what are the optimal imaging and stimulation parameters for minimizing opsin desensitization? We found that increasing the duration of two-photon imaging at 920 nm increases the rate of opsin desensitization, as evidenced by greater reductions in photostimulation responses across trials in stimulated neurons and the local network. This can be mitigated through a combination of longer inter-stimulus intervals and shorter imaging doses.

## 2. Materials and Methods

### 2.1 Animals, Viruses and surgery

Animal experimentation was carried out with approval from the UK Home Office and the University of Oxford Animal Welfare and Ethical Review Board. Mice were an even mixture of sexes (8 male, 8 females) with transgenic expression of GCaMP6s in excitatory neurons [mouse strains B6;DBATg(tetO-GCaMP6s)2Niell/J crossed with CamK2a-tTa(AI94)]. At 7 - 14 weeks old, mice were either implanted with a cranial window over the right somatosensory cortex (n=12) or right visual cortex (n=4). At the beginning of surgery, mice were anaesthetized with isoflurane (3%, 1 L/min) and kept under maintenance anesthesia for the rest of the procedure (1.5-2.5%, 0.3-0.5 L/min). Pre-operatively, Buprenorphine (Ceva) (1 unit/g), Meloxicam (Boehringer Ingelheim) (1 unit/g) and Dexamethasone (MSD) (5 units) were administered subcutaneously. The scalp was shaved and cleaned with an antiseptic solution of 2% chlorhexidine gluconate and 70% isopropyl alcohol (Becton Dickinson) before head-fixing the animal via ear bars on a stereotactic rig (Kopf instruments). Scalp and adipose tissue were excised from the right lateral side of the skull. A titanium headplate weighing ∼1 g with a 7 mm circular aperture was secured to the skull (-1.4 mm axial, +4 mm lateral / -1.9 mm axial, +3.8 mm lateral relative to bregma for somatosensory cortex implantation, +0.3 mm axial, +3.7 mm lateral relative to lambda for visual cortex implantation) using superglue and dental cement (SuperBond, Prestige Dental). For some mice, a second layer of 20% carbonated cement was used to further adhere the headplate to the skull. A 3/4 mm craniotomy was drilled with a dental drill (NSK) at the center of the headplate aperture and the dura was removed. A micropipette was front-loaded with pAAV-CamKIIa-C1V1(t/t)-mScarlet-KV2.1 (addgene) diluted 1:11 in sterile PBS. 500 nL of the virus was injected into the craniotomy 300 um deep from the cortical surface, at a rate of 50 nL/min using a hydraulic micromanipulator (Narashige). A cranial window comprised of two optically bonded glass coverslips (one the size of the craniotomy and one 1 mm larger) (Warner Instruments) was used to plug the craniotomy. The cranial window was secured in place using Cyanoacrylate (3M) and dental cement. The animal was allowed to fully recover for at least 3 weeks before any further experiments were conducted.

### 2.2 Two-photon Imaging and Stimulation

The microscope used in this study was a Bruker Ultima 2Pplus equipped with a tunable wavelength laser for imaging (Chameleon Ultra II, Coherent) and a 1035 nm fixed wavelength laser for photostimulation (Monaco 1035, Coherent). The 1035 nm laser was directed through a spatial light modulator (Neuralight 3D Ultra, Bruker). Photostimulation parameters and phase masks were defined and created using custom MATLAB software (NAPARM)^14^ before being loaded into external software. Imaging and photostimulation were controlled using Prairie View software (Bruker). Phase masks were loaded onto the SLM using BOSS (Boulder Nonlinear Systems).

The animal was head-fixed on a tip-tilt stage (TTR001, Thorlabs) under a 16x objective (N16XLWD-PF, Nikon). Stimulation and imaging triggers were coordinated using general purpose input/output software (PACKIO)^42^ running on a USB data acquisition card (National Instruments).

A reference two-photon image for opsin expression (conjugated to mScarlet) was captured at 765 nm using a 570-620 nm emission filter. NAPARM was used to group opsin-expressing neurons into groups of 10. Within each group, 6 mW 1035 nm laser light (2 MHz repetition rate) was simultaneously spiraled over each cell body. The spiral spanned a diameter of 10 um for a duration of 10 ms, repeating 25 times for a total photostimulation duration of 250 ms. The phase mask changed every 1 second and each group received 10 sets of spiral stimulations with a 15 s inter-trial interval. Photoresponsive neurons were identified manually from GCaMP6s ΔF/F stimulus triggered average images based on a baseline of 1 second pre-stimulus.

Photoresponsive cells were used for experimental testing. During the photoresponse decay experiments, each stimulation event consisted of spiraling 6 mW laser light over each cell body for 10 ms. The phase mask changed every 15 ms to target the other group of cells and each group received 10 blocks of stimulation in total (over 300 ms).

Responses to the photostimulation were imaged across a 1.4 x 1.4 mm field of view (1024 x 1024 resolution) at 15 Hz using resonant scanning at 920 nm (80 MHz pulse repetition rate, 50 mW average power after the objective), with a 500-550 nm emission filter.

### 2.3 GCaMP Imaging Analysis

Calcium imaging movies were processed using Suite2p^43^; cell bodies were automatically detected and carried forward for analysis. ΔF/F was computed for each cell body identified by Suite2p using the equation:

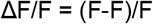

where

F = raw fluorescence

F = the mean of F in the 1 s pre-stimulation

Cells were labelled as ‘responsive’ in the photoresponse decay experiments if their mean ΔF/F in the 1 second post-stimulation was >= 0.1. The ‘response’ of a population of responsive cells to photostimulation is defined as the area under the curve of the mean ΔF/F across cells in the one-second post-stimulation.

If multiple cells were wholly or partially in the radius of a beamlet, only the cell with the centroid closest to the center of the beamlet was labelled as a ‘target’. When analyzing ‘non-target’ cells, any non-target cells wholly or partially within the radius of a beamlet were excluded from analysis.

For a cell to be labelled as excited or inhibited, it must be significantly correlated both across all trials (two-seconds around stimulation) concatenated and across trial one separately (p < 0.05 Pearson’s correlation). This was to ensure that correlated cells are active on trial 1 so that comparisons can be reliably drawn to the response on trial 10. The correlation coefficient of the cell was taken from the correlation across all trials

## 3. Results

### 3.1 Determining the baseline desensitization of optogenetically-evoked neuronal responses with varying 920 nm imaging duration

To assess optogenetic stimulus responses, we targeted sixteen putative cells in layer 2/3 somatosensory or visual cortex. The cells were split equally between two target groups using separate SLM phase masks (**Fig. 1(a)**). Ten trials were conducted to gather enough data to observe the effects of desensitization on neuronal responses. Cells with a centroid closest to the center of a beamlet (an individual point of light on the sample; see Methods) were labelled as ‘target’ cells if they were responsive within the radius of a beamlet. Initially, the stimulations were separated by an interval of 15 s as a common ISI used in all-optical experiments. The responses were collected by imaging using a 920 nm laser (1.4 x 1.4 mm field of view, 50 mW power on sample, 15 Hz, 1024 x 1024 pixels, ∼3 us total dwell time per cell; subthreshold dwell time per cell and power on sample for eliciting spikes^2,34^) and defined as the area under the curve of the mean ΔF/F across all target cells in the 1 s post-stimulation.

**Figure 1.**
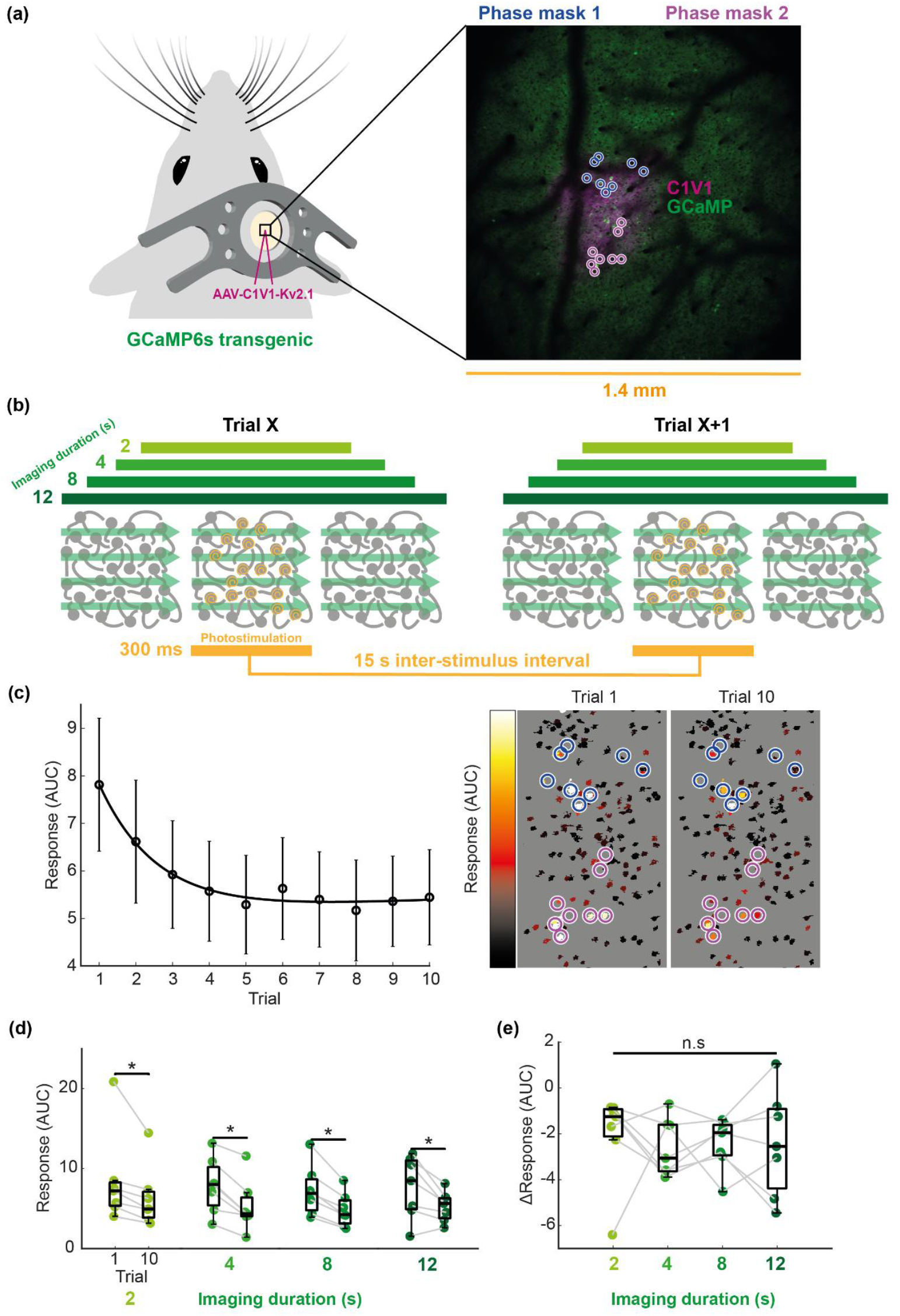
Determining the baseline desensitization of optogenetically-evoked neuronal responses with varying 920 nm imaging duration. The opsin AAV-C1V1-Kv2.1 was injected into layer 2/3 of either the right somatosensory cortex (n = 3) or right visual cortex (n = 4) of mice transgenic for GCaMP6s in excitatory neurons. Calcium activity was imaged over a 1.4 mm x 1.4 mm view around the opsin-expressing area (somatosensory cortex shown). Sixteen photostimulation targets were equally divided between two SLM phase masks. Schematic showing the different raster-scanning imaging durations centered around photostimulation with a fifteen second inter-stimulus interval. Each group of cells targeted by one SLM phase mask received ten blocks of 10 ms spiraling stimulation at 6 mW with a 20 ms interval between spirals. **(c)** Left: grand average response (over imaging conditions then animals, n = 7 mice) ±SEM fit with a two-term exponential model. Right: cells from an example session color-coded by their individual responses to stimulation with the same holograms as in **(a). (d)** Responses on trials 1 and 10 as a function of imaging dose (* = p < 0.05, Wilcoxon signed-rank test, n = 7 mice). Box plots show the median, upper quartile, lower quartile, maximum and minimum, excluding outliers (< lower quartile - 1.5 * inter-quartile range OR > upper quartile + 1.5 * inter-quartile range). Each dot represents an animal. Grey lines connect data from the same animal. **(e)** ΔResponse (trial 10 - trial 1 response) as a function of imaging dose (n.s = p > 0.05, Kruskal-Wallis test, n = 7 mice).

To investigate the impact of imaging with a 920 nm laser on C1V1 response, we varied the duration of imaging from 2-12 s centered on the photostimulation event (**Fig. 1(b)**). On average, the response of target cells decayed across trials, initially dropping by 20-30% in the first few trials and plateauing around trial five with responses remaining consistently at ∼70% of the initial response (**Fig. 1(c)**). Response desensitization over trials followed a similar trajectory across all imaging durations until trial five, when responses became comparatively less reduced for the 2 s imaging dose (**Fig. S1(a)**). Responses significantly decreased from trials one to ten, regardless of the imaging condition (**Fig. 1(d)**), with a trend towards larger imaging durations causing greater desensitization (**Fig. 1(e)**). To check that this was not an effect of the variation in initial pre-stimulation imaging duration, we confirmed that there were no significant differences in the first trial response between conditions (**Fig. S1(b)**). These results suggest that prolonged imaging doses at 920 nm may increase C1V1 desensitization.

### 3.2 Imaging at 920 nm increases desensitization at shorter inter-stimulus intervals

To further probe the effects of imaging at 920 nm on C1V1 desensitization under different experimental paradigms, we tested the two most extreme imaging doses: constant imaging and 2 s imaging, at inter-stimulus intervals (ISI) from 10-40 s (**Fig. 2(a)**). On average across all ISIs, both imaging conditions showed a significant decrease in responses from trials one to ten (**Fig 2(b)**), with the constant imaging condition causing significantly greater desensitization (**Fig. 2(c)**). The response decay curves for the two imaging conditions diverged as early as trial two (**Fig. 2(d)**).

**Figure 2.**
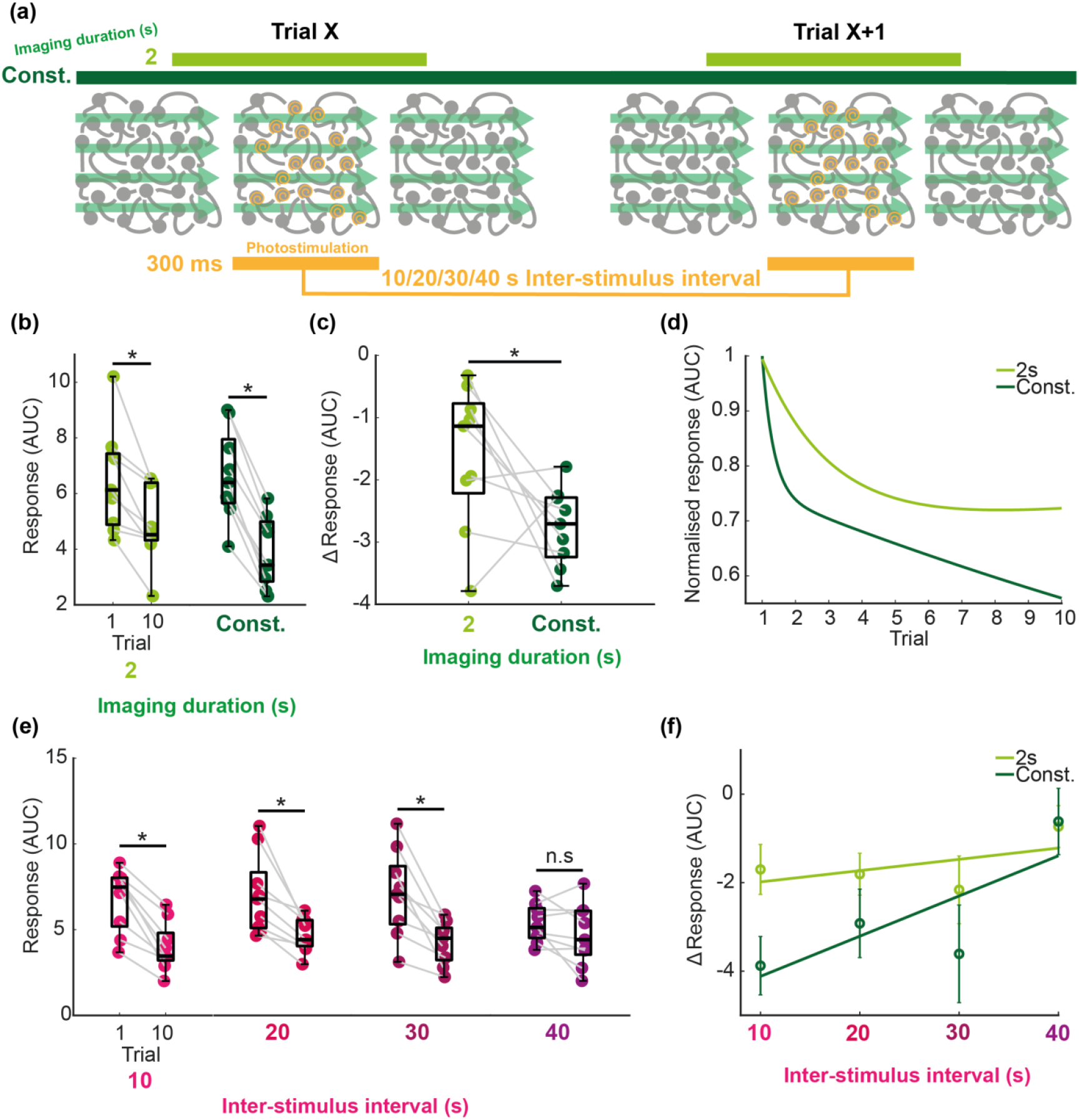
Imaging at 920 nm increases C1V1 desensitization at shorter inter-stimulus intervals. **(a)** Schematic showing the two imaging conditions (either imaging constantly or in a 2 s window centered around photostimulation) and four possible inter-stimulus intervals (ISI; 10/20/30/40 s). Each group of cells targeted by one SLM phase mask received ten blocks of 10 ms spiraling stimulation at 6 mW with a 20 ms interval between spirals. **(b)** Responses on trials 1 and 10 as a function of imaging dose, averaged across inter-stimulus intervals (* = p < 0.05, Wilcoxon signed-rank test, n = 9 mice). Box plots show the median, upper quartile, lower quartile, maximum and minimum, excluding outliers (< lower quartile - 1.5 * inter-quartile range OR > upper quartile + 1.5 * inter-quartile range). Each dot represents an animal. Grey lines connect data from the same animal. **(c)** ΔResponses (trial 10 - trial 1 response) as a function of imaging dose, averaged across inter-stimulus intervals (* = p < 0.05, Wilcoxon signed-rank test, n = 9 mice) **(d)** Normalized response (mean trial response averaged across inter-stimulus intervals then animals / mean trial 1 response averaged across inter-stimulus intervals then animals) as a function of trial number, fit with a two-term exponential model (n = 9 mice). **(e)** Responses on trials 1 and 10 as a function of inter-stimulus interval, averaged across imaging conditions (* = p < 0.05, Wilcoxon signed-rank test, n = 9 mice). **(f)** Average ΔResponse ±SEM fit with a first-degree polynomial (n = 9 mice).

On average across both imaging conditions, ISIs of 10, 20 and 30 s showed significantly decreased neuronal responses to stimulation from trials one to ten, but there was no significant desensitization for the 40 s ISI (**Fig. 2(e)**). Constant imaging exacerbated desensitization to a greater extent at shorter ISIs (**Fig. 2(f)**). These results demonstrate that desensitization of optogenetically-evoked neuronal signals increases as a function of imaging dose, but this relationship is only seen with ISI’s less than 40 s.

### 3.3 Imaging at 920 nm exacerbates desensitization of non-target cells across the field of view

It is possible that as targeted neurons respond less to stimulation, their ability to influence wider-scale network properties could change - a major confound for all-optical experiments. To investigate whether imaging at 920 nm had any effect on neuronal responses beyond the targeted cells, we assessed the activity of non-target cells across the field of view. Non-target cells were classified as either excited, inhibited or non-responders (**Fig. 3(a)**). For a cell to be labelled as excited or inhibited, its activity must be significantly correlated with at least one of the target cells (see Methods). Excited cells exhibited a different spatial distribution to Inhibited cells that was not dependent on opsin expression and was unaffected by imaging duration (**Fig. S2**). Like target cells, the responses of excited cells significantly decreased from trials one to ten, with the opposite being true for inhibited cells (**Fig. 3(b)**). Excited cells did not respond as strongly to stimulation as target cells and decayed in their responses much faster, plateauing by trial two only slightly above the non-responders (**Fig. 3(c)**). Larger imaging doses caused greater response desensitization across all inter-stimulus intervals (ISI) in excited cells (**Fig. 3(d)**; **Fig. S3**). Taken together, these results suggest that the amplitude of optogenetically-evoked responses in targeted cells non-linearly translates downstream, comparatively exacerbating the rate of response reduction in the local network. Thus, non-significant changes in the response of target cells, such as with a 40 s ISI, can cause dramatic changes in downstream responses. Therefore, during all-optical experiments, a combination of a short imaging dose and large ISI is needed to ameliorate response desensitization across the entire field of view (**Fig. 3(e)**), as a large ISI alone is not sufficient for non-target cells (**Fig. S3**).

**Figure 3.**
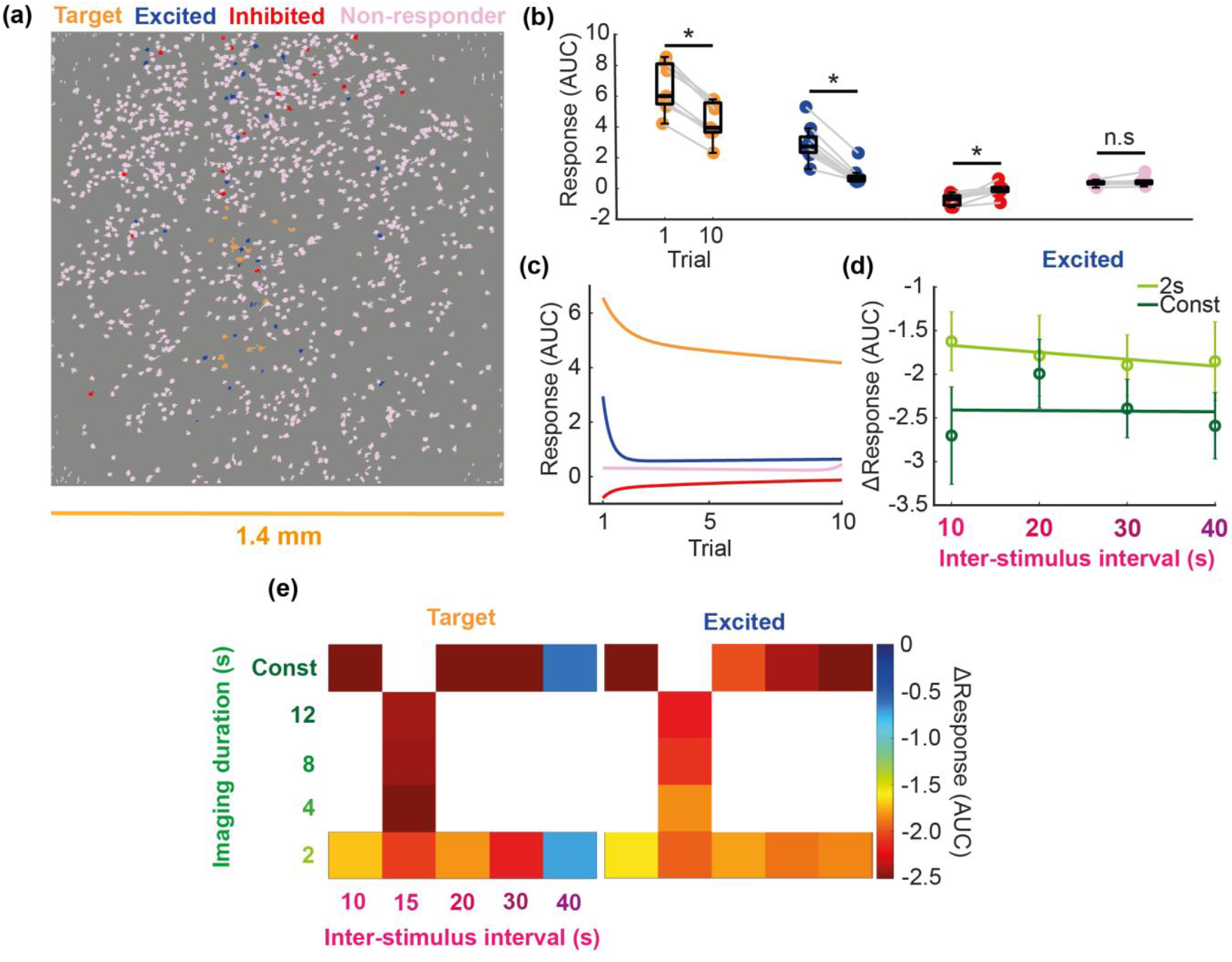
Imaging at 920 nm exacerbates C1V1 desensitization across the field of view. **(a)** Distribution of target, excited, inhibited and non-responder cells across the field of view from the same example session as in **Fig. 1(a,c). (b)** Responses on trials 1 and 10 for the different cell classifications (* = p < 0.05, Wilcoxon signed-rank test, n = 9 mice). Box plots show the median, upper quartile, lower quartile, maximum and minimum, excluding outliers (< lower quartile - 1.5 * inter-quartile range OR > upper quartile + 1.5 * inter-quartile range). Each dot represents an animal. Grey lines connect data from the same animal. **(c)** Grand average response (over conditions then animals, n = 9 mice) fit with a two-term exponential model. **(d)** Average ΔResponse (trial 10 - trial 1 response) ±SEM (n = 9 mice) of excited cells, fit with a first-degree polynomial. **(e)** ΔResponse in target and excited cells as a function of imaging dose and inter-stimulus interval. The same cells were stimulated across imaging conditions for the 10, 20, 30 and 40 s inter-stimulus intervals (n = 9 mice). For a different population of cells, the same cells were stimulated across imaging conditions for the 15 s inter-stimulus interval (n = 7 mice).

## 4. Discussion

The main advantage of all-optical interrogation over other less precise optogenetic stimulation methods, is dissecting causal relationships through the artificial recreation of endogenous-like cortical activity. Causality in systems neuroscience has been historically hard to define, with suggestions to instead broadly agree on methods to disambiguate causality, such as randomization, rather than dwell on definitions^44^. Whilst randomization is a conceptually sound gold standard to establish causality^45^, it has been difficult to achieve in practice^46^. In this regard, all-optical interrogation is uniquely positioned to determine whether specific neural activity or behavior is causally produced by other neural activity, or is just incidentally linked via a common input. However, recreation of endogenous-like activity requires careful calibration of photostimulation parameters such as laser power and scanning pattern^2^. As shown in this study, two-photon imaging at 920 nm can bias neural activity in C1V1-expressing neurons, throwing off the calibration. We also show that this bias can affect how injected activity is propagated and as such should also be considered when investigating causal relationships. Our results indicate that all-optical interrogation’s unique advantage is lost two-fold if opsin desensitization is not accounted for: firstly, if the inter-stimulus interval is not long enough, photostimulation-evoked activity in the targeted cells will decrease across trials, adding the influence of a confounding variable to any suggestions of causal relationships. Secondly, regardless of inter-stimulus interval, photostimulation-evoked activity in the surrounding network will decrease across trials, an effect that is exacerbated by larger imaging doses.

Since the first algal channelrhodopsin was discovered^3^, there has been a concerted effort to discover and engineer opsins with optimal kinetics for optogenetic modulation of neural circuits^36,47^. Such desirable traits include slower decay constants and larger photocurrents, which provide the high sensitivity preferred in two-photon stimulation experiments. Unfortunately, these characteristics also make opsins more vulnerable to cross-talk, which can complicate the interpretation of findings in all-optical studies. Typical approaches to reduce the impact of cross-talk between opsins and fluorescent reporters of neuronal activity in all-optical experiments include reducing the power of the imaging laser (which also reduces signal-to-noise) and scanning over large fields of view to reduce dwell time^17^. However, here we show that, over a large (1.4 mm x 1.4 mm) field of view, even low levels of opsin activation caused by imaging at 920 nm reduces photostimulation responses over trials, likely due to opsin desensitization. This is likely also true for other red-shifted opsins, as even those with peak excitation at higher wavelengths, such as ChrimsonR, still have spectral overlap with wavelengths used for GCaMP imaging^48–50^. With the aim of minimizing cross-talk, some studies have opted to image GCaMP at 1020 nm and stimulate channelrhodopsin-2 at 920 nm^51^. However, while channelrhodopsin-2 is not efficiently excited at 1020 nm, small photocurrents are still produced^39^ which will likely cause similar problems to those seen in the current study. Many all-optical studies use a combination of a red-shifted opsin, a green fluorescent reporter, constant two-photon imaging and inter-stimulus intervals of <=10 s^10,11,13,14,16,41^. Under these conditions, if the identity of the targeted cells remained constant over successive stimulations, there would be a significant decrease in the photostimulation-evoked responses of both the targets and downstream excited cells across trials. Minimizing response reduction over successive stimulations across the entire field of view requires combining a long inter-stimulus interval (>=40 s) with intermittent short imaging doses.

In this study, we focus on how reductions in photostimulation-evoked responses can be mitigated in traditional all-optical experiments using red-shifted opsins and green fluorescent reporters of neuronal activity. However, changing the opsin/sensor combination can also be used to mitigate opsin desensitization via cross-talk. For example, one might use a spectrally-distinct combination of blue-shifted opsins with red calcium sensors^35,52^. The downside here is that red calcium sensors are generally less sensitive with slower kinetics^53^. Another option is using red-shifted step-function opsins (SFO) in combination with green calcium sensors. SFO’s can be opened or closed with different wavelengths of light. Therefore, to prevent subthreshold activation of the opsin during imaging outside of the stimulation, one can simultaneously scan with the wavelength that negatively regulates opsin activity. For example, a previous study identified a channelrhodopsin-2 variant (sdChR) that is activated by blue-light, but only in the absence of orange light^48^. The downside here is that incorporation of an SFO into a two-photon all-optical experiment would either require the integration of a third wavelength laser into the light path, the use of a fast-switching programmable tunable wavelength laser for positive/negative stimulation, or an SFO that is negatively-modulated by a one-photon source with a wavelength that is outside of the emission filter bounds of the photomultiplier tube.

In conclusion, our results demonstrate that a combination of optogenetic inter-stimulus intervals >=40 s and a 2 s imaging dose around each stimulation significantly mitigates desensitization of targeted C1V1-expressing cells and ameliorates desensitization of excited cells across the field of view during all-optical interrogation. This will reduce an external source of trial-by-trial variability in future all-optical experiments. The extent to which imaging will impact opsin desensitization in other all-optical experiments will depend on laser wavelengths, power on sample, dwell time per cell (combination of duty cycle, zoom, resolution and soma size), spectral overlap of the opsin and reporter, and opsin expression per cell. Therefore, the experiments described herein will have to be repeated for unique all-optical paradigms to find the best tradeoff between data quantity and opsin desensitization. In the absence of such data, one’s best option will be to minimize the imaging dose and increase the inter-stimulus interval as much as is practical.

## Data availability

The data generated for this study will be publicly shared upon peer-review publication.

## Code availability

The computer code written for this study will be publicly shared upon peer-review publication.

## Disclosures

The authors declare no conflicts of interest related to the research, authorship, or publication of this article.

## Acknowledgments

This research was supported by funding from the Wellcome Trust (204651/Z/16/Z) to A.M.P and an ERC grant (funded by UKRI, EP/X026655/1) to A.L.

## Author information

### Contributions

B.R. and A.M.P. designed the study. R.M.L. provided two-photon microscopy training and implanted cranial windows in three of the animals. B.R. conducted all other surgeries and all experiments. B.R. performed analysis with advice from A.M.P. A.M.P and A.L. provided supervision. All authors wrote the Manuscript.

## Notes

### Competing Interest Statement

The authors have declared no competing interest.

