## Supplementary figures for "Desensitization of opsin responses during all-optical interrogation depends on imaging parameters"

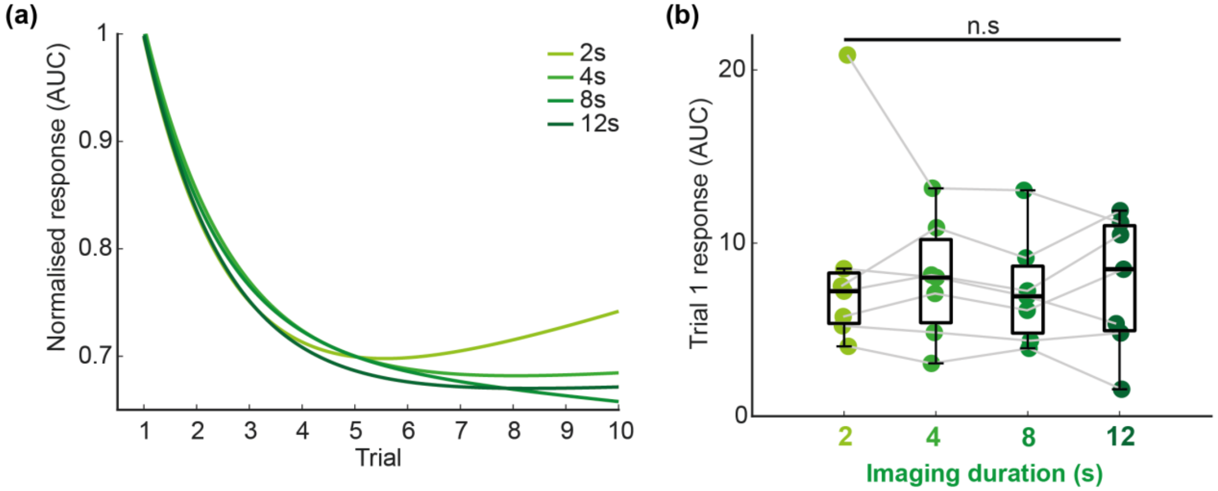


**Figure S1 | C1V1 desensitization dynamics over trials under different imaging conditions**
**(a)** Normalized response (mean trial response across animals / mean trial 1 response across animals) as a function of trial number (n = 7 mice). **(b)** Responses on trial 1 as a function of imaging dose (n.s = p > 0.05, Kruskal-Wallis test, n = 7 mice). Box plots show the median, upper quartile, lower quartile, maximum and minimum, excluding outliers (< lower quartile - 1.5 * inter-quartile range OR > upper quartile + 1.5 * inter-quartile range). Each dot represents an animal. Grey lines connect data from the same animal.

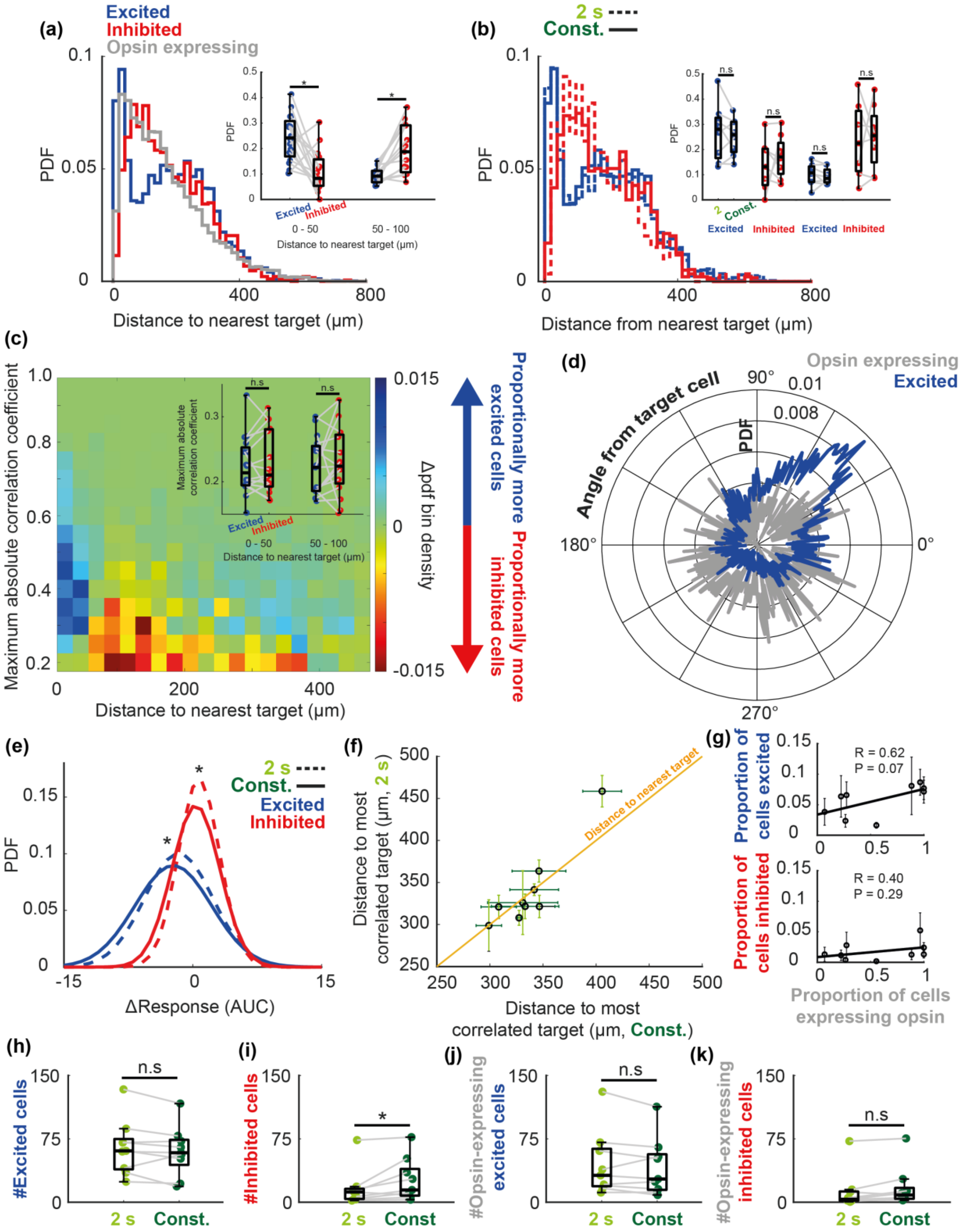


**Figure S2 | Spatial distribution of excited and inhibited cells under different imaging conditions**
**(a)** Probability distribution of excited, inhibited and opsin-expressing non-target cells as a function of distance from the nearest target cell. Inset: Each point is data from an individual animal, averaged

across all cells across all conditions (* = $p < 0.05$, Wilcoxon signed-rank test, n = 16 mice). Box plots show the median, upper quartile, lower quartile, maximum and minimum, excluding outliers (< lower quartile - 1.5 * inter-quartile range OR > upper quartile + 1.5 * inter-quartile range). Grey lines connect data from the same animal. **(b)** same as in **(a)** but separated by imaging condition. Inset: (* = $p < 0.05$, Wilcoxon signed-rank test, n = 9 mice). **(c)** Maximum correlation strength of an individual non-target cell with any target as a function of distance from the nearest target cell, color-coded by excited bin count - inhibited bin count. Inset: Each point is data from an individual animal, averaged across all cells for a given type across all conditions (* = $p < 0.05$, Wilcoxon signed-rank test, n = 16 mice). **(d)** Polar plot of an example mouse with opsin expression confined to a part of the field of view. The polar plot shows the probability distribution for excited and opsin-expressing cells at different angles from a target cell, averaged across targets across all conditions. The probability distributions for excited and opsin-expressing cells are non-overlapping **(e)** probability distribution of excited and inhibited cells under different imaging conditions (* = $p < 0.05$, two-sample Kolmogorov-Smirnov test, pooled across n = 9 mice). **(f)** Mean ±SEM distance of non-target cells from their most correlated target cell under different imaging conditions. Orange line is a representation of what a 1:1 relationship would look like, for example if distance to nearest target was plotted on each axis instead, as this would not vary between imaging conditions. **(g)** Correlation between the proportion of cells in the field of view that are excited/inhibited and the proportion of cells expressing opsin (R = Pearson correlation coefficient). **(h-k)** The number of different classifications of non-responsive cells under different imaging conditions (* = $p < 0.05$, Wilcoxon signed-rank test, n = 9 mice).

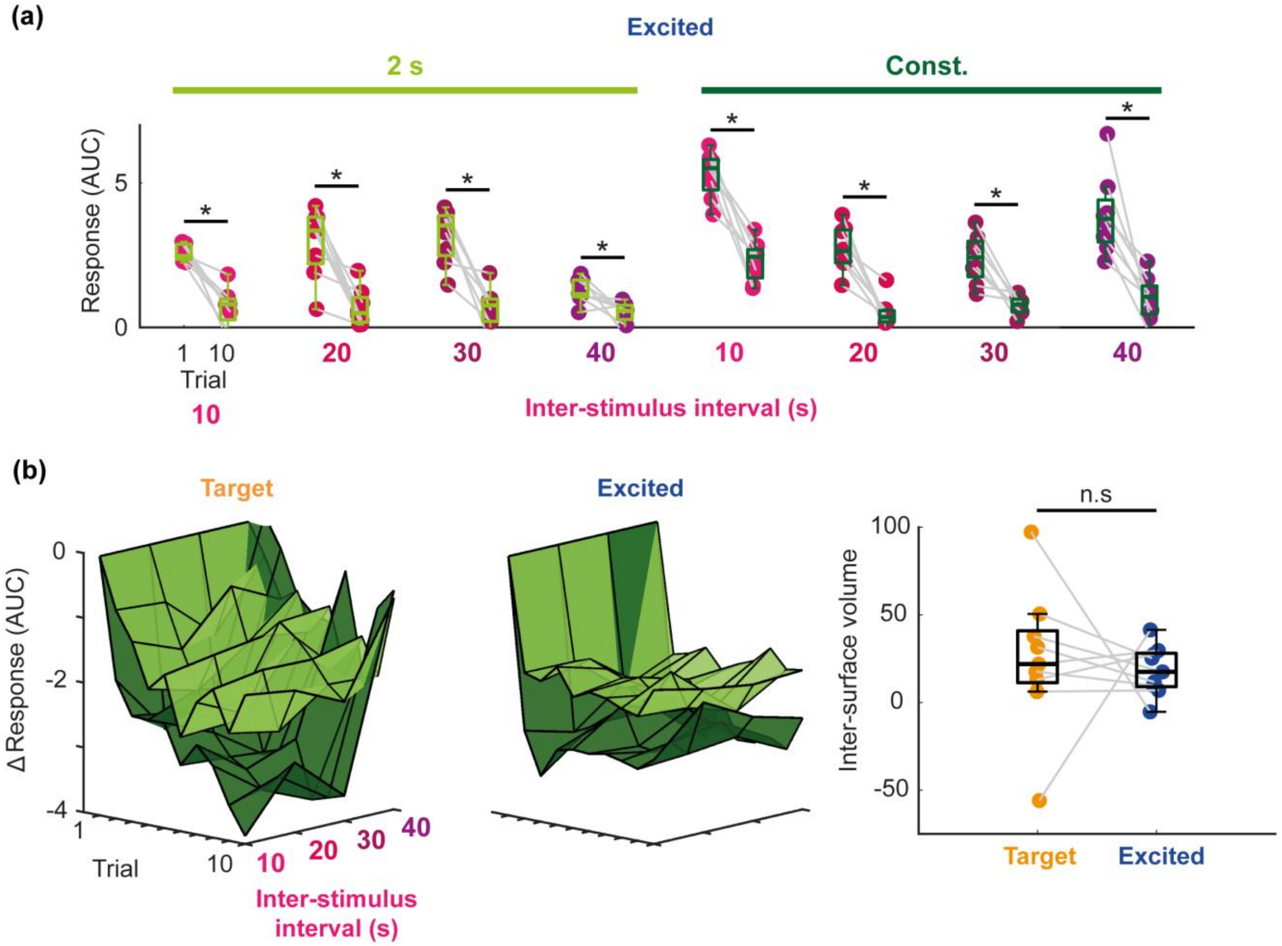


**Figure S3 | Desensitization of excited cell responses cannot be rescued by a 40 s inter-stimulus interval**

**(a)** Excited cell responses on trials 1 and 10 as a function of inter-stimulus interval under different imaging conditions (* = $p < 0.05$, Wilcoxon signed-rank test, $n = 9$). Box plots show the median, upper quartile, lower quartile, maximum and minimum, excluding outliers (< lower quartile - 1.5 * inter-quartile range OR > upper quartile + 1.5 * inter-quartile range). Each dot represents an animal. Grey lines connect data from the same animal. **(b)** Surface plots showing target cell (left) and excited cell (middle) ΔResponses (trial response - trial 1 response) across trials and inter-stimulus intervals under different imaging conditions. Right: Volume differences between the 2 s surface plot and constant imaging surface plot for target and excited cells (area under 2 s imaging surface - area under constant imaging surface. * = $p < 0.05$, Wilcoxon signed-rank test, $n = 9$).
